# Effective infection with dengue virus in experimental neonate and adult mice through intranasal route

**DOI:** 10.1101/2022.02.18.481117

**Authors:** Minyue Qiu, Lixin Zhao, Junjie Zhang, Yalan Wang, Minchi Liu, Dong Hua, Xiaoyan Ding, Xiaoyang Zhou, Jie Zeng, Huacheng Yan, Jintao Li

## Abstract

As a member of the family Flaviviridae, the dengue virus is transmitted to humans through mosquito vectors. The virus causes dengue fever with flu-like symptoms, life-threatening dengue hemorrhagic fever, and dengue shock syndrome that affects four billion people in 128 countries. It can also be transmitted through atypical routes, such as needle stick injury, vertical transmission, and the receipt of contaminated blood or organs. Although sporadic cases cannot be classified as atypical transmission routes and raise respiratory exposure concerns, the risks remain unclear. Here, we analyzed the respiratory infectivity of the dengue virus in BALB/c suckling and A6 adult mice using the dengue virus serotype 2 and intranasal inoculation. The results showed that the clinical symptoms of intranasally infected mice included excitement, emaciation, malaise, and death, and histopathological changes and viremia dynamics were observed in multiple tissues, including the brain, liver, and spleen. Notably, the DENV serotype 2 showed clear brain tropism after intranasal infection, and viral loads peaked at seven days post-inoculation. Furthermore, the virus was isolated from mouse brains, and the sequence homology between the viral stock and dengue virus isolates was 99.99%. Similar results were observed in adult IFN-α/β receptor-deficient mice. Taken together, our results suggest that the DENV serotype 2 can infect suckling mice and adult immune-deficient mice via the nasal route. The results of our study broaden our perception of atypical dengue transmission routes and may help explain dengue virus infections that occur without the presence of mosquito vectors.

**Importance:** Dengue is transmitted mainly via mosquito vectors. Flu-like dengue fever or life-threatening severe dengue threatens four billion people in 128 countries. Non-vector dengue cases have occurred occasionally since 1990. These cases cannot be classified as atypical transmission routes and are suspected to have arisen via sexual or respiratory transmission. Recently, the sexual transmission of dengue has been confirmed in humans. The respiratory infectivity of other members of the Flaviviridae family, such as Zika and Japanese encephalitis viruses, has been determined under experimental conditions. The potential for respiratory infection with the dengue virus has not yet been confirmed. In this study, we analyzed the respiratory infectivity of the dengue virus in suckling mice and adult immunodeficient mice via intranasal inoculation, thereby exploring the potential for respiratory transmission in cases with non-vector dengue infection, especially for patients who were exposed to dengue virus contaminants.

## Introduction

The seasonal emergence of the dengue virus (DENV) has raised global concerns due to the expansion of areas affected by the epidemic, consequently increasing the risk of infection (1, 2). DENV is a member of the family Flaviviridae and is transmitted mainly through mosquito bites. The pathogen causes self-limiting flu-like dengue fever (DF) or life-threatening severe dengue, such as dengue hemorrhagic fever (DHF) and dengue shock syndrome (DSS) (3). In the last 50 years, the morbidity associated with DENV infections has increased dramatically, threatening four billion people in 128 countries (4). Notably, in addition to the primary transmission route, cases caused by atypical transmission routes such as needle stick injury, vertical transmission, and receipt of contaminated blood or organs have occasionally been documented since 1990 (5-10). Sporadic cases that occur in the absence of the vector in pandemic areas also include the possibility of respiratory transmission of DENV (11).

Recently, it was reported that other members of the family Flaviviridae, such as Zika and Japanese encephalitis virus (JEV), can infect guinea pigs or mice via intranasal inoculation (12, 13). Previous studies have shown similar results for the West Nile virus, St. Louis encephalitis virus, and hepatitis C virus (HCV) (14-16). However, the possibility of intranasal infection with DENV has not been confirmed or verified, which is intriguing because 35%–38% of DENV patients show respiratory symptoms including sore throat, cough, and nasal congestion (17). In addition, although rare, the virus has been isolated from cerebrospinal fluid, saliva, urine, and from the upper respiratory tract of patients with respiratory symptoms (18-21).

To determine whether DENV can infect experimental animals through the intranasal route, BALB/c suckling mice, which are widely used in DENV research (22, 23), and adult IFN-α/β receptor-deficient mice (A6 mice), which are susceptible to DENV (24), were challenged with dengue virus serotype 2 (DENV-2) by intranasal inoculation. In this study, we identified the intranasal transmission route of DENV under experimental conditions for the first time, and the results suggest that DENV may have the potential for respiratory transmission. These results may also facilitate a better understanding of non-vector-borne disease outbreaks in DENV pandemic areas.

## Results

### Suckling mice can be efficiently infected by DENV-2 via the intranasal route

We used suckling mice, which are widely used for virus fixation, isolation, virulence detection, and virulence recovery in DENV-related research, to confirm the possibility of intranasal transmission of DENV (Figure 1A). As an intracranial challenge results in infection in suckling mice, we compared infectivity via intranasal DENV inoculation relative to that in intracranially inoculated suckling mice. Three-day-old suckling mice were inoculated intranasally with 2.4×10^4^ PFU DENV-2, and the weight, clinical signs, and vital status of each animal were monitored thereafter. We noticed that the onset of infection symptoms usually occurred at 7 to 9 days post-inoculation (dpi), which was several days later than in the intracranially challenged group, and all challenged animals showed the same clinical symptoms, including emaciation and malaise at later stages (Figure 1B). However, partially intranasally challenged animals also showed excitement in the early stages (Supplementary Table 1), along with mania and sensitivity to moving objects. Infections in both groups resulted in 100% mortality; however, the mean survival time of suckling mice that were inoculated intranasally with viral stock was delayed from 6 to 9 days using the same viral titer administered to mice with intracranial infection (Figure 1C). Further, we evaluated the dose-dependent survival rates for suckling mice in the intranasally challenged group; mortality occurred in 100%, 62.5%, and 33.3% of the animals inoculated with 2.4×10^4^, 2.4×10^3^, and 2.4×10^2^ PFU of DENV-2, respectively (Figure 1D). Therefore, we selected a titer of 2.4×10^4^ PFU for subsequent experiments. In addition, the process of infection was consistent with the resulting weight changes in the animals. In the early stages, the weight change rates of mice in the challenge groups nearly coincided with those in the control group. However, once infection onset occurred, the weight change rates decreased significantly in mice in the challenged groups regardless of the infection stage, and most animals died after the weight change rate decreased to 100% or less (Supplementary Figure 1).

**Figure 1.**
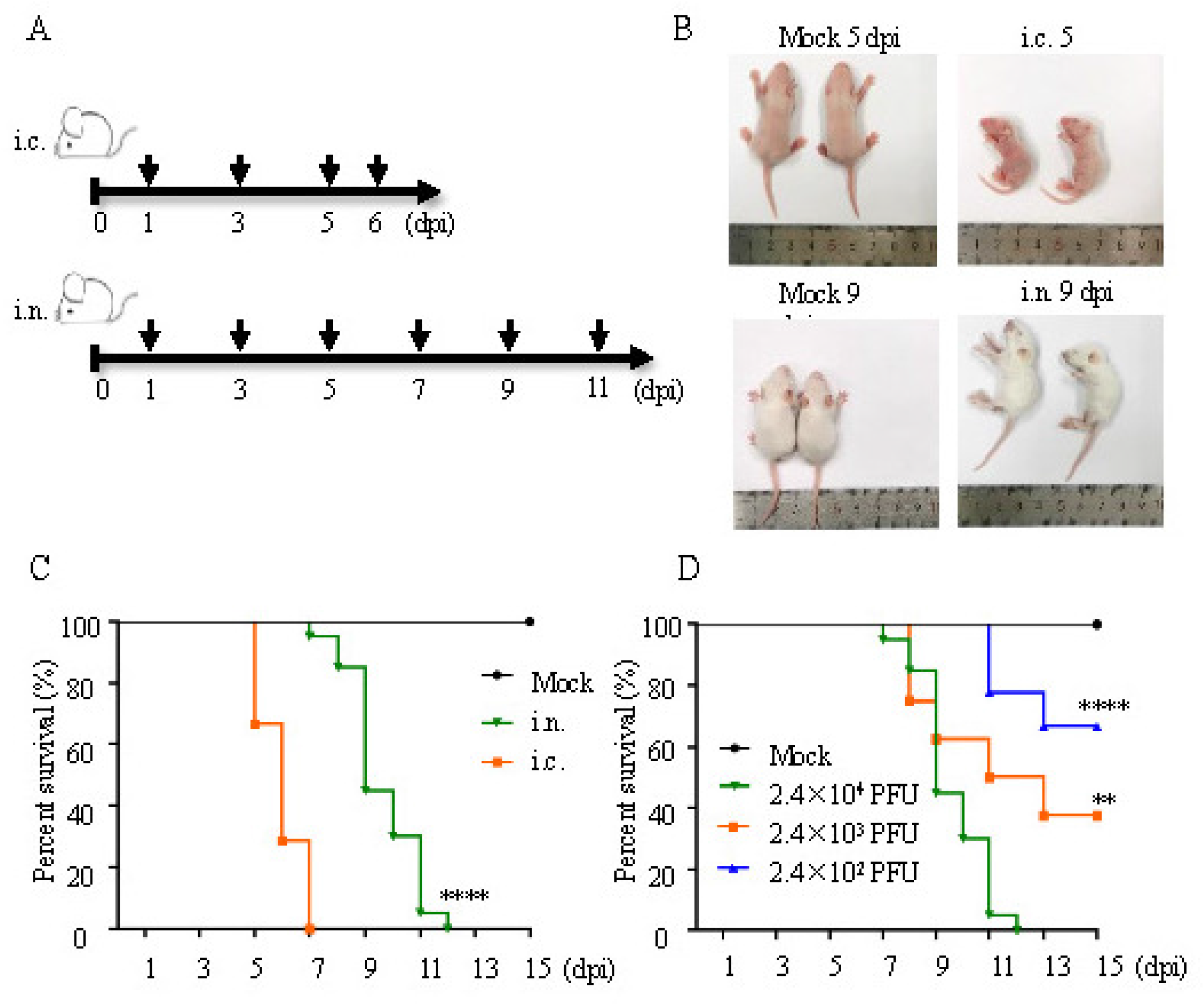
DENV-2 infection via the intranasal route is lethal in BALB/c suckling mice. (A) Experimental scheme. On the third day after birth, the newborn BALB/c suckling mice were challenged intracranially or intranasally with DENV-2. The body weight, clinical phenotypes, and mortality rates were evaluated, and organs (heart, liver, spleen, lung, kidney, and brain) and the blood of challenged suckling mice were harvested (black arrow). (B) Representative images of intracranially or intranasally treated BALB/c suckling mice. (C) Survival proportion of DENV-2 treated suckling mice infected via intracranial or intranasal routes. Survival conditions were monitored daily after challenge (titer = 2.4×10^4^ PFU; i.c., n = 21; i.n., n = 20; mock, n = 8). (D) Survival proportion of DENV 2 intranasally infected suckling mice with different titers.

### The brain is the main target of DENV-2 after intranasal inoculation

The heart, liver, spleen, lungs, kidneys, brain, and blood of intranasally inoculated suckling mice were harvested at 1, 3, 5, 7, 9, and 11 dpi to detect the viral load. Owing to the shorter mean survival time, the tissues of animals in the intracranially challenged group were harvested at 1, 3, 5, and 6 dpi. In the intranasally challenged group, viremia was detected in all animals at 5 dpi by qRT-PCR, with peak loads of 4.55 log (copies)/mL (Figure 2A). Additionally, DENV-2 RNA was detected in multiple organs at 5 dpi by qRT-PCR, with peak loads ranging from 3.47 to 3.88 log (copies)/g (Supplementary Figures 2A-E). In brain tissues, DENV-2 RNA was detected at 3 dpi, and the virus replicated linearly, reaching a peak with an average viral load of 7.8 log (copies)/g at 7 dpi (Figure 2B), which was approximately twice as high as in the other organs. Viremia and higher viral loads were observed 2 days earlier in the intracranially challenged mice than in intranasally challenged animals, (Supplementary Figures2A-E). Although DENV-2 replicated more rapidly in the brain tissues of intracranially challenged mice, resulting in an early peak in viral load (Figure 2B), the difference in the final RNA copy number between the two groups was not significant (Supplementary Figure 2F). In addition, immunofluorescence staining showed clear fluorescence signals in the brains of intranasally inoculated suckling mice at 9 dpi, when the clinical phenotype related to DENV-2 infection was most obvious (Figure 2C). The results of electron microscopy analysis confirmed the presence of DENV-2 particles in the brain of suckling mice at 9 dpi (Figure 2D).

**Figure 2.**
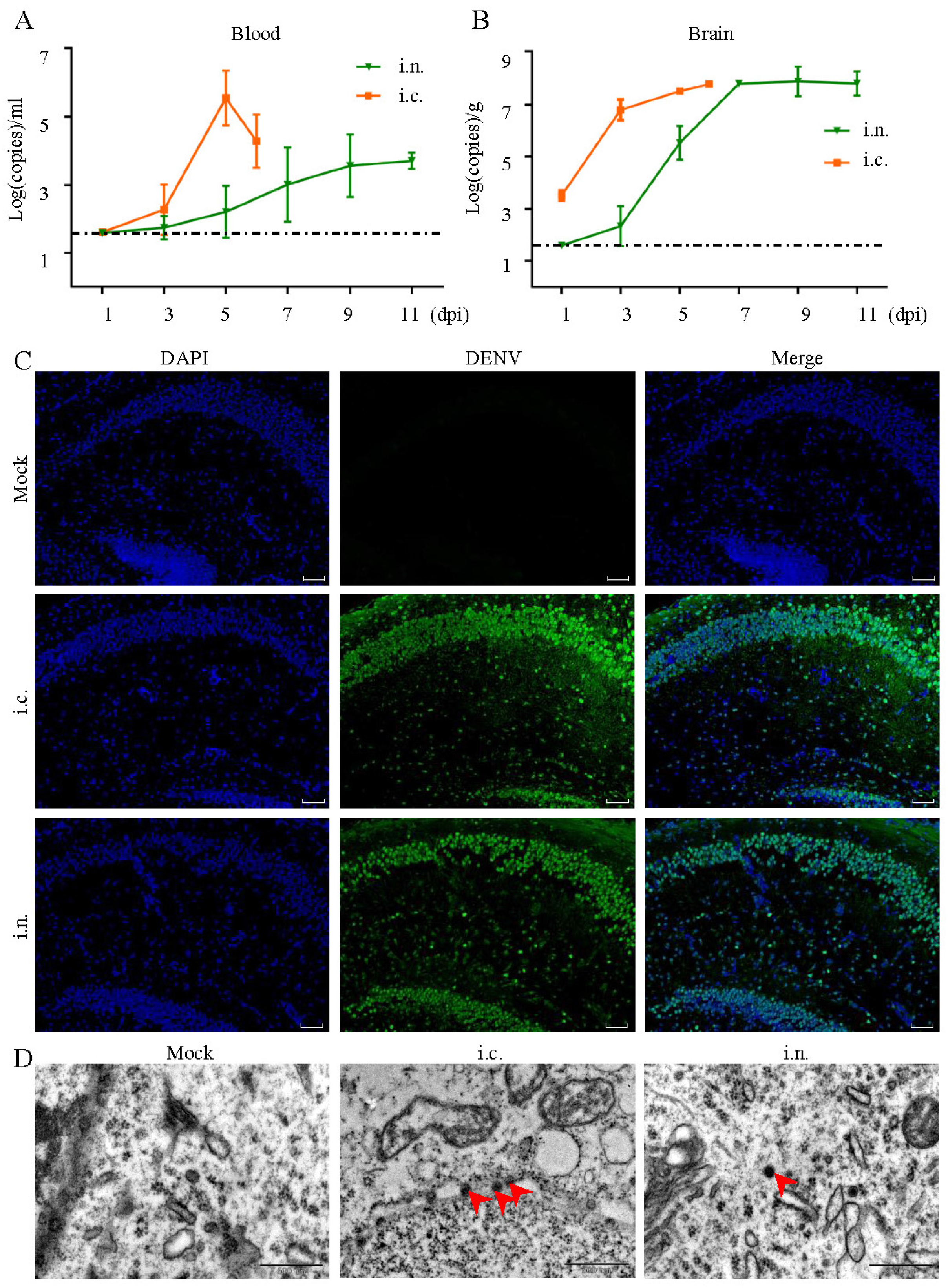
Infection status of DENV-2 treated suckling mice. A. Blood samples were harvested, and viremia was evaluated (each time point, n = 4 or n = 5). The dotted black line indicates the limits of detection. B. Brain samples were harvested, and the virus titer was evaluated by RT-PCR (each time point, n = 4 or n = 5). C. Immunostaining of the hippocampus from intracranially and intranasally challenged animals at the severe illness phase (5 and 9 dpi, respectively). Scale bar: 50 μm. D. Representative transmission electron micrographs showing DENV particles in DENV-2 challenged suckling mouse brains.

### Histopathological changes in multiple organs of mice after intranasal inoculation

Next, we characterized the pathological changes that occurred in the challenged suckling mice. Despite the lower viral loads in organs besides the brain, intranasal infection with DENV-2 resulted in substantial pathological changes in the liver and spleen at late infection stages. The histopathological phenotypes were similar to those in intracranially challenged animals and included modest inflammatory cell infiltration and a few multinucleated giant cells in the liver, as well as numerous apoptotic bodies and inflammatory cell infiltration in the spleen (Figure 3A). No obvious histopathological changes were observed in other organs (Supplementary Figure 3). Histopathological changes in the brain tissue were observed in the hippocampus and cerebral cortex in both the intracranially and intranasally challenged mice. A large number of necrotic pyramidal cells with pyknotic, broken, or dissolved nuclei, eosinophilic cytoplasm, and modest gliocyte proliferation were observed in all areas of the hippocampus in the intracranially challenged mice. Moderate pyramidal cell necrosis was observed in the CA2, CA3, and DG areas of the hippocampus in the intranasally challenged mice. The main histopathological changes in the cerebral cortex included neuronal necrosis and vacuolization, and these phenotypes were modest in the intracranially challenged mice, whereas the phenotypes were extensive in the intranasally challenged mice (Figure 3B).

**Figure 3.**
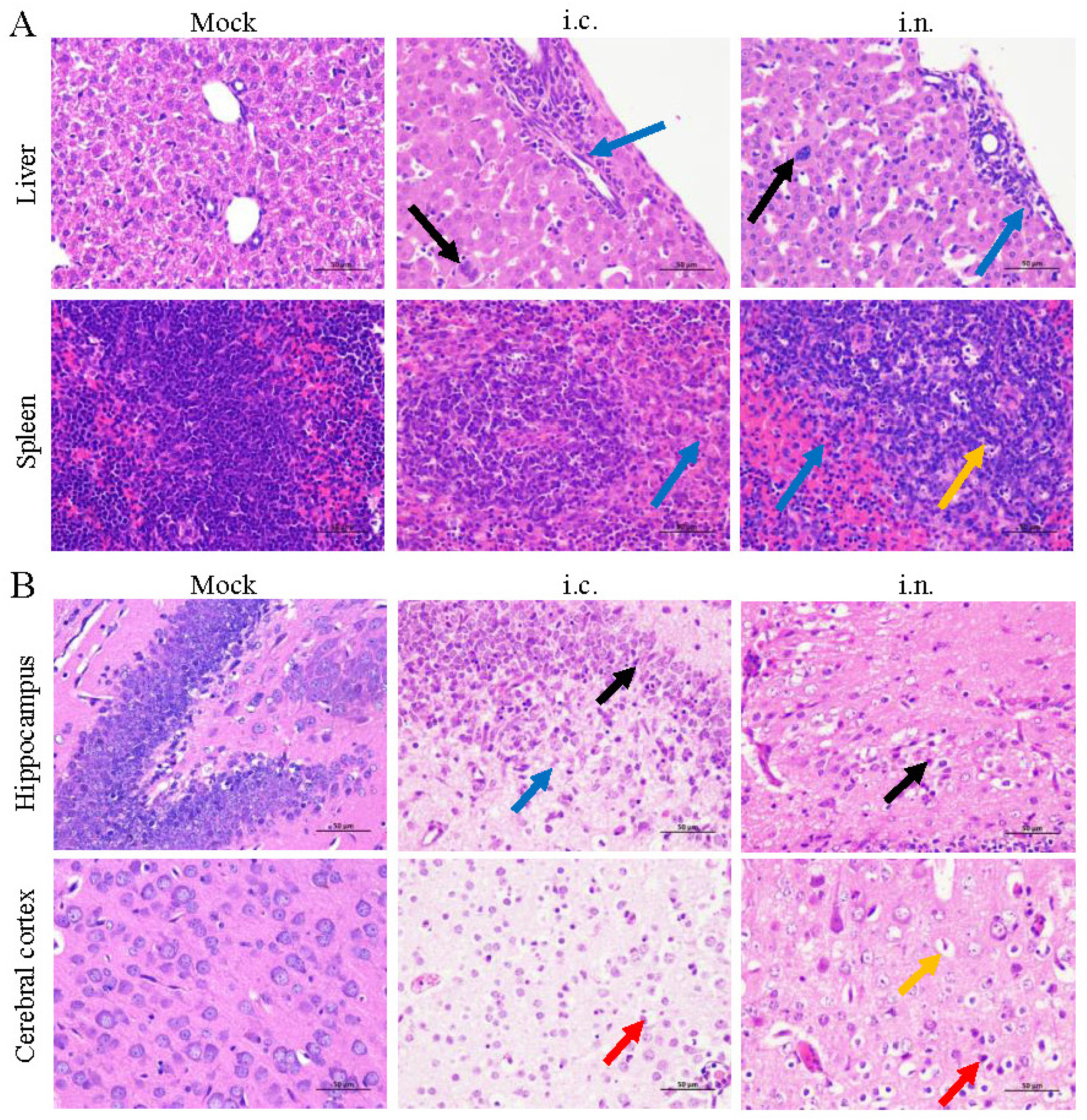
DENV-2 infection results in pathological changes. A. Selected organs were harvested from intracranially or intranasally challenged mice sacrificed at a late phase of illness (5 and 9 dpi, respectively) and were processed for hematoxylin and eosin (H&E) staining. Blue arrows, granulocytes; black arrows, multinucleated giant cells; orange arrows, apoptotic bodies. B. Brains were harvested from intracranially or intranasally challenged mice sacrificed at a late phase of illness (5 and 9 dpi, respectively) and processed for H&E staining.

### DENV-2 can be isolated from intranasally inoculated mice and efficiently replicates in Vero cells

After intracranial or intranasal infection, moribund mice were sacrificed and anatomized to harvest the brain tissue which showed the highest viral load. Both DENV-2 challenged mouse groups showed growth and plaque morphology similar to the viral stock in the supernatant of ground brain tissue (Figure 4A). Next, the virus in the supernatant of the ground brain was serially passaged three times. Cytopathic effects, viral RNA copy numbers, and infectious virion numbers were evaluated at each passage. The results showed that DENV-2 isolates from the two challenged mouse groups could induce cytopathic effects (CPE) in Vero cells within 5 days after inoculation (Supplementary Table 2). DENV-2 from intracranially and intranasally challenged animals replicated effectively in Vero cells, resulting in high viral RNA loads (Figure 4B). However, as shown in Figure 4C DENV-2 isolated from intranasally challenged animals assembled and released greater numbers of mature virions after replication in Vero cells compared to intracranially challenged mice. Furthermore, the sequence homology between the viral stock and DENV-2 isolates was 99.99%, and the DENV-2 isolate showed 99.68% sequence homology with the DENV-2 New Guinea C strain (GenBank: KM204118.1) (Supplementary Figure 4).

**Figure 4.**
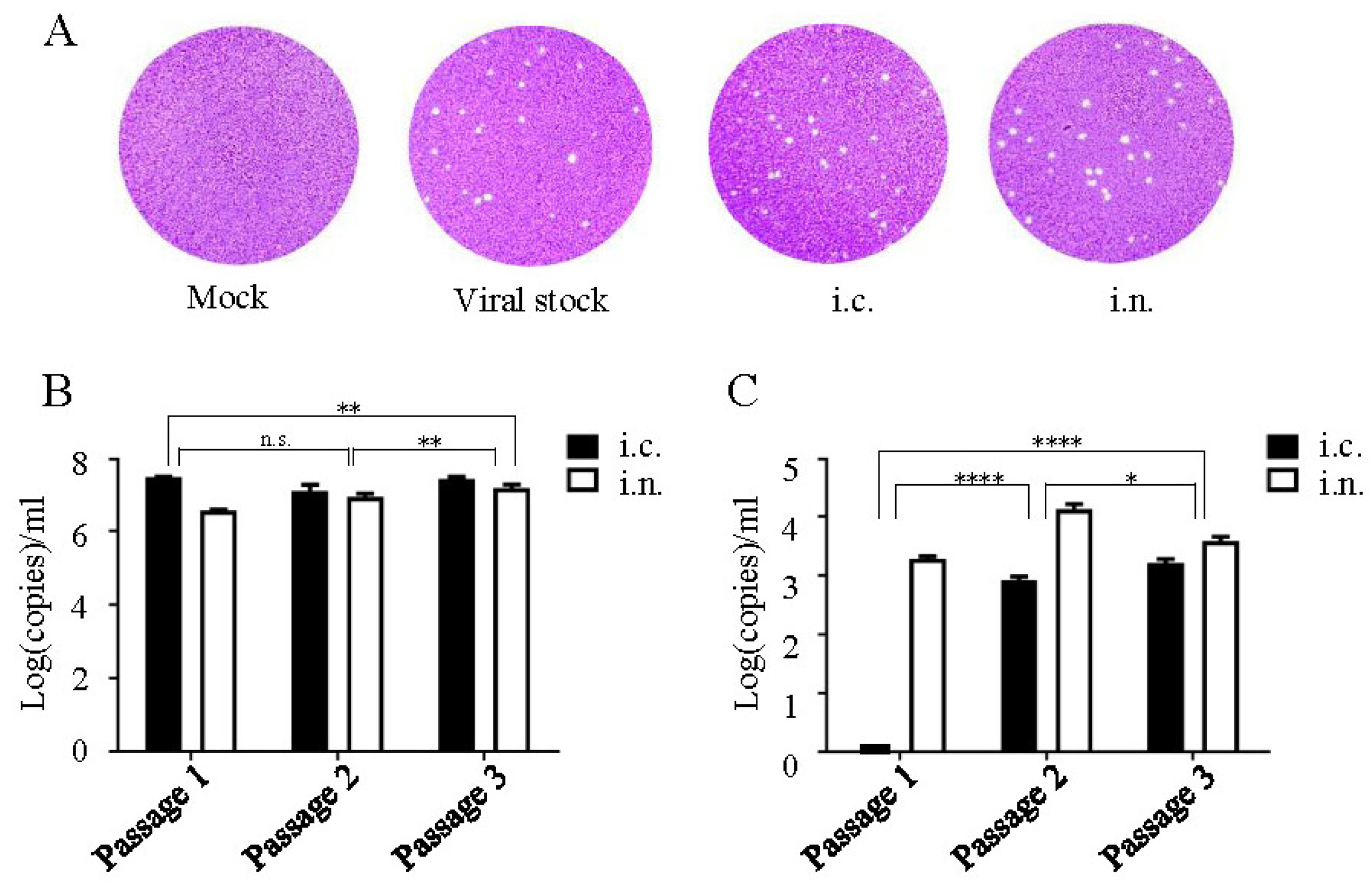
Isolation of DENV-2 from Vero cells. A. Plaque morphology of DENV-2 viral stock and isolates from intracranially or intranasally infected mouse brains. B. DENV-2 was isolated from Vero cells and cultured for three passages, and the viral load was evaluated via RT-qPCR. C. The viral titer during each passage was evaluated in Vero cells.

### Adult A6 mice can be efficiently infected by DENV-2 via the intranasal route

To validate the intranasal infectivity of DENV-2 in adult mice, groups of 4–6 weeks old male A6 mice were challenged with 2.4×10^5^ PFU DENV-2. The weight, clinical signs, and vital status of each animal were recorded. The results showed that A6 mice could be infected with DENV-2 intranasally. The clinical symptoms of intranasally infected A6 mice included emaciation, malaise, hind leg paralysis (Figure 5A), and death. At 7 dpi, 100% of the animals exhibited weight loss (Supplementary Figure 5A), and 50% of the animals were dead at 11–12 dpi (Figure 5B), with weight change rates lower than – 30%. Interestingly, 50% of intranasally challenged animals showed recovery from neurological symptoms at 9–11 dpi, with mice showing regained body weight (Supplementary Figure 5A). Moreover, after inoculation with higher viral titers, dose-dependent survival rates were observed (Supplementary Figure 5B). A study of moribund animals showed that intranasal infection caused mild and transient viremia in mice at 3–5 dpi (Figure 5C), indicating poor nose-to-blood migration of DENV-2 in adult mice. Robust viral replication and proinflammatory factor production in the mouse brain (Figure 5D-E) indicated that the brain rather than the respiratory system may serve as the preferred target of DENV-2 infection after intranasal infection.

**Figure 5.**
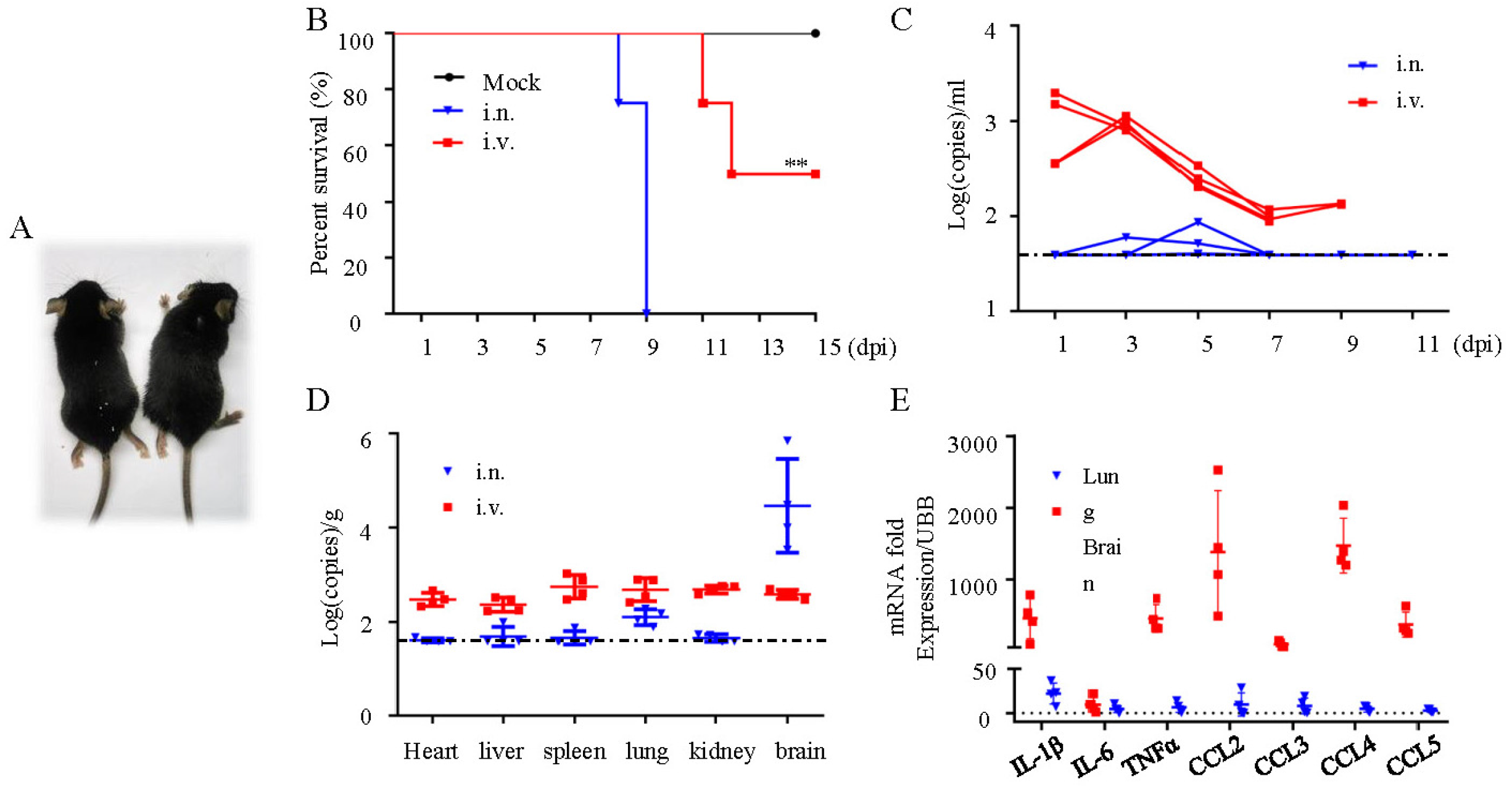
DENV-2 infection via the intranasal route is lethal in A6 mice. (A) Representative images of intranasally treated A6 mice with neurological symptoms. (B) Survival proportion of DENV-2 treated A6 mice administered the virus via the intravenous or intranasal routes. Survival conditions were monitored daily after challenge (titer = 2.4×10^5^ PFU; i.c., n = 4; i.n., n = 4; mock, n = 4). (C) Viremia in A6 mice after DENV-2 inoculation. (i.v., n = 4; i.n., n = 4). Dotted lines indicate the limits of detection. (D) Internal organ samples were harvested, and the virus titer was evaluated (i.v., n = 4; i.n., n = 4). (E) Expression of pro-inflammatory factors in the brains and lungs of moribund A6 mice.

## Discussion

The re-emergence of DENV is a major public health concern that has put people in the tropical and subtropical regions at risk. In the 1950s, *Aedes* mosquitoes were shown to play an important role in the transmission of DENV (25). Subsequently, several atypical transmission routes for DENV were characterized in humans, such as needlestick injury, vertical transmission, and receipt of contaminated blood or organs. However, rare cases (11, 18) of suspected respiratory exposure to DENV in the absence of a mosquito vector were not stressed because of the lack of experimental confirmation and criticism concerning the failure of mosquito prevention (11, 26). In this study, we found that DENV-2 can infect both wild-type suckling mice and adult immunodeficient mice administered intranasally under experimental conditions, with dose-dependent infection outcomes. Because DENV infects and replicates in the human primary lung epithelium and lung cancer cell lines (27), and has been isolated from the upper respiratory tract of patients (18, 19), our findings further strengthen the evidence for DENV infectivity in the upper respiratory mucosa *in vivo*.

After intranasal inoculation, infected animals showed systemic infection and neurological symptoms, such as excitement/hind leg paralysis, emaciation, malaise, and death. Viremia was detected in all animals and substantial histopathological changes were observed in multiple tissues. However, the brain was the main target of DENV-2 after intranasal inoculation, rather than the lungs, and the occurrence may be related to different infection mechanisms. As part of the immune system, the nasal mucosa can contain inhaled antigens (28). After the intrusion and replication of the virus in nasal epithelial cells, monocytes may differentiate into macrophages, which are reportedly the target cells for DENV (29). Subsequently, DENV-carrying macrophages may reflux into the blood and cause systemic infections. Because DENV-2 RNA copies in the brain were higher and detectable earlier than in other tissues and blood, the results suggest that the respiratory tract plays an important role in early DENV-2 infection. After DENV-2 invades the upper respiratory tract mucosa, DENV-2 may enter the brain directly through the olfactory nerve, leading to massive viral replication and various neurological symptoms. Subsequently, the recruited leukocytes may increase the severity of mouse encephalitis owing to the Trojan horse effect (30, 31). We noted that the morbidity and mortality of murine models varied after challenge with different titers of DENV-2, and animals challenged with the same titer exhibited varying degrees of infection. This result indicates that individual differences exist in the infection of DENV through nasal routes, and hosts are infected by DENV only when their nasal mucosa is exposed to an infective titer of the virus. In addition to respiratory exposure, a new non-vector transmission route, i.e. sexual transmission of DENV, has been suspected (32). Until recently, there have been isolated case reports describing the sexual transmission of DENV in humans from non-pandemic areas (33). This result indicates that the natural mucous membrane-dependent transmission route is feasible but rarely recognized. There are several potential explanations for this finding. First, viable virus titers in body fluids other than the blood may exist in a few cases, and these require closer examination (18). Second, the ubiquity of mosquito vectors in epidemic areas (34) makes it difficult to exclude the possibility of a primary transmission route in doubtful cases. In addition, the 3–14 day incubation period of DENV and the poorly understood respiratory transmission feature makes epidemiological investigation more difficult (35). Importantly, people may have ignored the potential of alternate transmission routes for DENV, such as intranasal, aerosol, and sexual transmission routes, which may explain why the bulk of atypical transmission cases occur where mosquito prevention practices are rigorously implemented, such as in hospitals and laboratories, or in non-endemic areas (36-38).

This study had several limitations. First, the mechanism of the upper respiratory tract to brain infection remains unclear. The blood-brain barrier (BBB) is a complex structure that constitutes a barrier between the blood and the central nervous system (39). BBB destruction caused by DENV infection may lead to encephalopathy or encephalitis, which is associated with morbidity in 5%–7% of patients and mortality in ∼50% of patients (40). Further investigation of the mechanisms underlying upper respiratory tract-to-brain infection will help understand the mechanisms of DENV-associated encephalopathy or encephalitis. Other nasal routes, such as inhalation of DENV droplets or aerosols, should be analyzed in animal models to fully understand the potential for respiratory transmission of DENV. Finally, the infectious titers of DENV in this study were suitable for animal models alone. Whether DENV can be transmitted in human populations through the nasal route remains unknown. As described above, it is difficult to accurately identify the cause of infection in individuals living in pandemic areas. As DENV prevention and control programs have progressed (41, 42), there may be more non-vector dengue cases in the future.

In conclusion, this study showed that DENV-2 can cause acute infection both in suckling mice and adult A6 mice via intranasal inoculation, resulting in neurological symptoms. Upper respiratory tract cells may be the first station for viruses to invade and replicate, and the mechanisms underlying the upper respiratory tract-to-brain infection require further investigation. These findings indicate that the respiratory mucosa of mice can be intruded by DENV-2 and cause further infection with sufficient titers, suggesting that the potential for respiratory transmission of DENV may be greater than assumed previously. The possibility of inhalation or contact with contagious droplets or aerosols should be considered for cases of non-vector DENV infection, especially for healthcare workers who may be exposed to concentrated DENV contaminants.

## Methods

### Cells and viruses

Vero cells were cultured for 5−8 passages in DMEM medium (Gibco, USA) with 10% fetal bovine serum (FBS), and incubated in a humidified atmosphere containing 5% CO^2^ at 37°C. Cells at 80%−85% confluency were used for inoculation with DENV-2. DENV-2 was cryopreserved and provided by the Center for Disease Control and Prevention of the Southern Military Theatre. Three-day-old BALB/c suckling mice were inoculated with DENV-2 via the intracranial route, and the brain was harvested at 5 dpi and quickly ground in PBS on ice. The ground tissue was collected and centrifuged at 10,000 ×*g* for 10 min. The supernatant was collected gently and centrifuged at 10,000 ×*g* for 10 min. The procedure was repeated twice. The collected supernatants were stored as viral stocks at −80°C. Titers were evaluated using a plaque assay on Vero cells.

### Animal infection

Healthy 30-35g pregnant BALB/c mice were purchased from the Animal Center of the Army Medical University (Third Military Medical University, Chongqing, China) and provided sterile water and chow ad libitum, and acclimatized for 5-7 days prior to baby mouse birth. After giving birth, 3-day-old suckling mice were inoculated intranasally with 2.4×10^5^−2.4×10^2^ PFU DENV-2 using 10-fold serial dilutions ofPBS. The blood of infected mice was sampled daily. The challenged animals were monitored for clinical phenotypes, morbidity, and mortality. Clinical phenotypes were evaluated independently by three observers in a single-blinded manner. Moribund mice were euthanized with isoflurane and anatomized to harvest the heart, liver, spleen, lungs, kidneys, and brain. The tissue samples were fixed in 4% paraformaldehyde for 24 h at 4°C, embedded in paraffin, and sectioned. To determine the viral kinetics, tissues and blood were harvested every other day from 1 dpi onwards and ground in a homogenizer with precooled TRIzol (Invitrogen, USA). Tissues were quickly homogenized on ice for RNA isolation. Animals in the positive control group were inoculated intracranially with 2.4×10^5^ PFU DENV-2. Animals in the negative control group were inoculated intranasally with 3 μL supernatant from healthy ground suckling mouse brains. A6 mice were provided by the Department of Microbiology, School of Basic Medical Sciences, Capital Medical University, China. Male mice aged 6–8 weeks were challenged intranasally with 2.4×10^5^–2.4×10^2^ PFU DENV-2 using 10-fold serial dilutions of PBS, and the primary inoculation viral titers were determined based on the body weight of suckling mice and adult A6 mice. The experiments were conducted as previously described. Animals in the positive control group were inoculated intraperitoneally with 2.4×10^5^ PFU DENV-2, and the animals in the negative control group were inoculated intranasally with 30 μL supernatant from healthy ground suckling mouse brains.

### RNA isolation and qRT-PCR

Total RNA was extracted from the heart, liver, spleen, lungs, kidneys, brain, and blood using TRIzol reagent (Tiangen Biotech, China) according to the manufacturer’s recommendations. The PrimeScript™ RT Reagent Kit (TaKaRa, Japan) was used to reverse-transcribe cDNA from total RNA. Quantitative real-time reverse transcription-PCR (qRT-PCR) was performed using the TB Green Premix Ex Taq II (TaKaRa, Japan) with the LightCycler® 96 system (Roche, USA). Serial dilutions of the DEMV-2 plasmid with known concentrations were used to establish the standard curve and evaluate the amplification efficiency of the system. RNA copies per mL or RNA copies per gram of each sample were calculated from the Cq values using quantitative PCR based on analysis using the standard curve. The relative expression levels of IL-1β, IL-6, TNFα, CCL2, CCL3, CCL4, and CCL5 were determined via RT-PCR. The primers used in this study are listed in Supplementary Table 3.

### Immunofluorescence staining

Paraffin sections of the mouse brain were dewaxed using graded xylol and hydrated with graded ethanol. The sections were incubated with 3% hydrogen peroxide solution for 25 min to reduce endogenous peroxidase activity. Sections were boiled in 0.01 M citrate buffer (pH 6.0) for 10 min to repair the antigens. Normal goat serum (Abcam, England) was used to block non-specific staining for 2 h at 37°C. Next, sections were incubated with anti-dengue virus 1+2+3+4 primary antibodies (ab26837, Abcam, England) at 4°C overnight, and were subsequently incubated with FITC-conjugated secondary antibodies (Servicebio, China) for 2 h. The sections were stained with 4,6-diamidino-2-phenylindole (DAPI) (ab228549, Abcam, England) for 4 min. Representative sections were scanned using a Pannoramic DESK digital slice scanner (3D HISTECH, Hungary) with the Pannoramic Scanner software.

### Histopathology staining

Mouse tissue paraffin sections were dewaxed with xylol, hydrated with ethyl alcohol, and stained with hematoxylin for 3 min and eosin for 5 min. Images were photographed using a 40× objective with the Eclipse Ci-L microscope (Nikon, Japan) equipped with a DS-U3 camera (Nikon, Japan).

### Transmission electron microscopy

Brain tissues were quickly trimmed into cuboids with a volume of ∼1×1×2 mm^3^. The cuboids were fixed with 2.5% glutaraldehyde overnight at 4°C and washed with PBS for 15 min three times. Then, the cuboid was fixed again with 2% osmic acid for 2 h. Following graded acetone dehydration, tissues were infiltrated, embedded, and polymerized in resin. The resin mass was sliced to a thickness of 70 nm, and sections were stained with uranyl acetate and lead citrate and imaged using a JEM 1200EX (JEOL, Japan) transmission electron microscope.

### Virus isolation and identification

Moribund suckling mice were sacrificed and anatomized to harvest their brains at 5 or 9 dpi (intracranially or intranasally inoculated mice, respectively). The brain was quickly ground with PBS, centrifuged, and the supernatant was obtained. After cell confluency reached 80%, Vero cells were transfected with a 100-fold diluted brain supernatant and maintained in DMEM medium (Gibco, USA) supplemented with 2% FBS (Gibco, USA) at 37°C in 5% CO_2_. After seven days, the culture media and cells were collected and this constituted passage #1. Additional passages were performed using the same method with 10-fold dilutions of the early generation viral stock. Viral RNA was detected by qRT-PCR, as described above, and viral titers were evaluated using plaque assays.

### DENV-2 genome sequencing and analysis

Total virus RNA was extracted from DENV-2 or mock infected mice brain using the Purelink RNA minikit (Life Technology). Next, cDNA was produced by using Moloney murine leukemia virus (Mo-MLV) reverse transcriptase (TaKaRa, Japan) with a reverse primer, and 15primer pairs were used to generate overlapping amplicons spanning the entire genome accordingly. All sequencing was performed using the ABI 3730xl sequencer (ABI, USA). The genome sequences of DENV-2 were assembled by mapping the reads to the reference genome (Accession No. KM204118.1) of the DENV-2 New Guinea C strain using the DNA STAR v7.1 software.

### Plaque assay

DENV-2 isolates were serially diluted with DMEM medium (Gibco, USA) from 10^−2^ to 10^−6^ cells and added to 6-well plates with Vero cell monolayers of 80-85% constancy. The plates were incubated for 2 h at 37°C. After the supernatant was removed carefully, 2 ml of semisolid maintenance medium (1.2% methylcellulose, 2% FBS in DMEM medium) was added to each well and incubated for 5–7 days at 37°C. Subsequently, the semi-solid medium was removed carefully, and 1% crystal violet (Beyotime, China) was added for staining for 20–30 min. The plaques were counted after rinsing and air drying.

### Ethics statement

All animal and cell experiments were performed under biosafety level 2 conditions and conformed to the Chinese Regulations of Laboratory Animals. All experimental procedures were approved (AMU20170525) by the Ethics Committee for Animal Experimentation of the Army Medical University and followed the National Institutes of Health Guide for the Care and Use of Laboratory Animals.

### Data analysis

All data were analyzed using GraphPad Prism v6.01 software. Data are shown as the mean ± SD. The log-rank test was used for survival analysis. Two-way analysis of variance (ANOVA) followed by Tukey’s test was used to evaluate the significance between passages and viral origins. In all tests, values of *P* < 0.05 were considered significant.

## Acknowledgments

This work was supported by National Natural Science Foundation of China (grant number 81570497) and Natural Science Foundation of Chongqing (grant number cstc2020jcyj-bshX0116). We would like to thank Prof. Jing An (Department of Microbiology, School of Basic Medical Sciences, Capital Medical University) for generously providing the A6 mice.

